# Palmitoyl transferase ZDHHC20 promotes pancreatic cancer metastasis

**DOI:** 10.1101/2023.02.08.527637

**Authors:** Goran Tomić, Clare Sheridan, Alice Y. Refermat, Marc P. Baggelaar, James Sipthorp, Bhuvana Sudarshan, Cory A. Ocasio, Alejandro Suárez-Bonnet, Simon L. Priestnall, Eleanor Herbert, Edward W. Tate, Julian Downward

## Abstract

Metastasis is one of the defining features of pancreatic ductal adenocarcinoma (PDAC) that contributes to poor prognosis. In this study, the palmitoyl transferase ZDHHC20 was identified in an *in vivo* shRNA screen as critical for metastatic outgrowth, with no effect on proliferation and migration *in vitro*, or primary PDAC growth in mice. This phenotype is abrogated in immunocompromised animals, and in animals with depleted natural killer (NK) cells, indicating that ZDHHC20 affects the interaction of tumour cells and the innate immune system. Using a chemical genetics platform for ZDHHC20-specific substrate profiling, a number of novel substrates of this enzyme were identified. These results describe a role for palmitoylation in enabling distant metastasis that could not have been detected using *in vitro* screening approaches and identify potential effectors through which ZDHHC20 promotes metastasis of PDAC.

## Introduction

Metastasis, a spread of cancer cells from the primary tumour to secondary sites within the body, is frequently present in patients with pancreatic cancer. Approximately 3 out of 5 patients have distant metastases at the time of PDAC diagnosis (Strobel et al., 2019). In patients who undergo resection and adjuvant chemotherapy, the disease recurs in 65% of cases in the form of local recurrence and distant metastasis (Jones et al., 2019). The majority of distant metastases occur within two years, with half of liver metastases being detected within 12 months after surgery. In addition, there is evidence to suggest that early dissemination of tumour cells is a feature of PDAC pathogenesis (Rhim et al., 2012).

The *KRAS* gene is frequently mutated in human PDAC and the mutation is considered to be the key initiating event in pancreatic cancer (Ying et al., 2016). Other frequently mutated genes in pancreatic cancer are associated with G1/S checkpoint, TGF-ß pathway, histone modifiers, SWI/SNF complex, BRCA pathway, and WNT signalling (Bailey et al., 2016). The metastatic potential of these tumours is linked with mutant *TP53, SMAD4* (*DPC4*) and TGF-ß pathway, as well as oncogenic *KRAS* gene dosage (Iacobuzio-Donahue et al., 2009; Morton et al., 2010; Mueller et al., 2018; Whittle et al., 2015; Zhong et al., 2017). However, the contribution of other PDAC-associated genes to secondary site seeding is not fully explored. *In vitro* genetic screens are an important tool to interrogate the contribution of specific genes to cellular phenotypes, although not well suited for mimicking the metastatic cascade. *In vivo* genetic screens can provide more relevant answers as cancer cells can interact with the microenvironment (Chen et al., 2015). An additional level of relevance is added when these screens are carried out in an immunocompetent context, enabling investigation of cancer cell dependencies not detectable in other settings.

We aimed to investigate the *in vivo* metastatic potential of genes associated with pancreatic cancer. Here we report and characterise the palmitoyl transferase ZDHHC20 as a novel critical mediator of PDAC metastasis, which has no effect on primary tumour growth, but is necessary for the metastatic expansion. The expression of ZDHHC20 in tumour cells renders them resistant to NK cell-mediated cytotoxicity, allowing distant metastasis to the liver and lung. Substrate profiling identified a number of targets of ZDHHC20, whose palmitoylation could affect metastatic ability of tumour cells.

## Results

### ShRNA screen identifies mediators of PDAC metastasis *in vivo*

We carried out an *in vivo* shRNA screen in a pooled format using the TRC shRNA backbone (Figure 1A) to investigate the role of pancreatic cancer-associated genes in metastatic outgrowth. This platform has previously been used for effective shRNA screening under both *in vitro* and *in vivo* conditions (Huang et al., 2012; Miller et al., 2013), and our selection of candidate genes included those with mutations detected in large-scale pancreatic cancer sequencing studies (Biankin et al., 2012; http://cancer.sanger.ac.uk/cosmic; Jones et al., 2008). Given that *Kras* is a critical driver of pancreatic cancer, its downstream effectors may play a role in promoting metastasis, and a number of studies have looked at downstream gene expression to gain insight into the cellular changes implemented by this oncogene (Arena et al., 2007; Horsch et al., 2009; Loboda et al., 2010; Mukhopadhyay et al., 2011; Zhou et al., 2010). We compared *Ras* gene signatures with genes that show differential expression between normal and cancerous pancreatic samples in Oncomine (Rhodes et al., 2004), allowing us to derive a more selective *Ras* signature with increased relevance to pancreatic cancer. The set of candidate genes (284) was selected using three criteria: association with pancreatic cancer, frequency of mutation in pancreatic cancer, or regulation by *Ras* (Figure S1, Supplementary Table 1). Cells used in the screen were derived from a liver metastasis in the *Kras*^*G12D/+*^;*Trp53*^*R172H/+*^;*Pdx1-Cre* (KPC) genetically engineered mouse model of pancreatic cancer.

**Figure 1.**
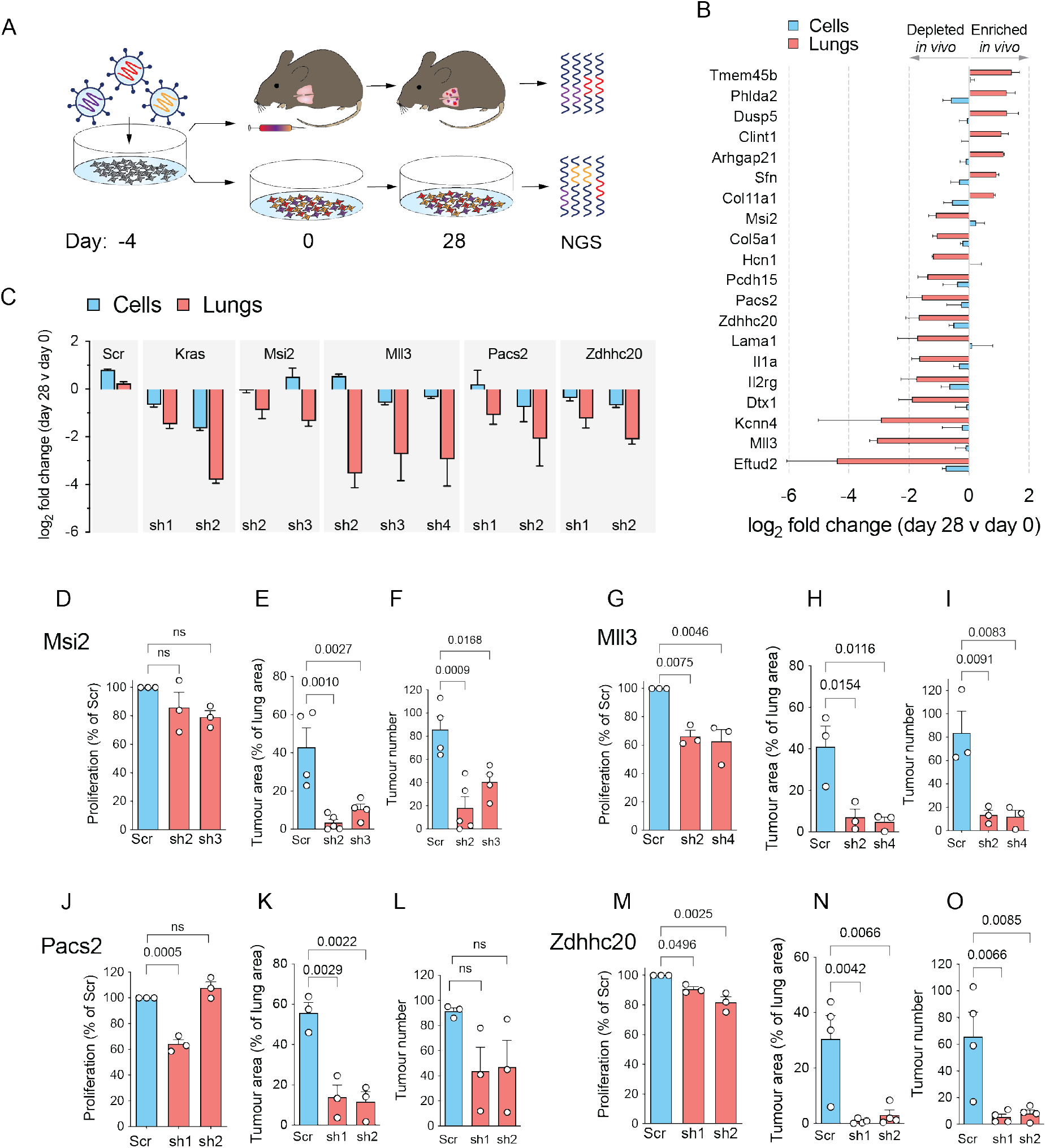
*In vivo* shRNA screen identifies novel mediators of pancreatic cancer metastasis. **(A)** Schematic of the shRNA screen. **(B)** Average log_2_ fold change of multiple shRNAs per gene *in vivo* (lungs) versus *in vitro* (cells). **(C)** Individual shRNA representation of scrambled (Scr) and selected shRNAs *in vivo*. **(D-F)** Proliferation in 2D of cells expressing scramble or shRNAs targeting *Msi2* **(D)**, quantification of tumour area **(E)** and tumour number **(F)** in the lungs of mice injected with the scramble shRNA or two shRNAs targeting *Msi2* cells after 4 weeks. **(G-I)** Proliferation in 2D of cells expressing scramble or shRNAs targeting *Mll3* **(G)**, quantification of tumour area **(H)** and tumour number **(I)** in the lungs of mice injected with the scramble shRNA or two shRNAs targeting *Mll3* cells after 4 weeks. **(J-L)** Proliferation in 2D of cells expressing scramble or shRNAs targeting *Pacs2* **(J)**, quantification of tumour area **(K)** and tumour number **(L)** in the lungs of mice injected with the scramble shRNA or two shRNAs targeting *Pacs2* cells after 4 weeks. **(M-O)** Proliferation in 2D of cells expressing scramble or shRNAs targeting *Zdhhc20* **(M)**, quantification of tumour area **(N)** and tumour number **(O)** in the lungs of mice injected with the scramble shRNA or two shRNAs targeting *Zdhhc20* cells after 4 weeks. Data are shown as mean + S.E.M. Ordinary one-way ANOVA with Tukey’s multiple comparisons test was used for data in **D-O**. P-values are shown above the comparison lines, except for non-significant differences, where ns is indicated instead.

To focus on the later stages of metastasis, colonisation of the secondary tissue and outgrowth of macrometastasis, intravenous injection was used to deliver tumour cells into the bloodstream and thereby observe distant organ seeding. We compared shRNA abundance prior to injection of cells (Day 0) and in lung tumours at day 28 post-injection by next generation sequencing. Enrichment of an shRNA indicated that ablation of gene expression provides a selective advantage for tumour cells and implies that the target gene is a metastasis inhibitor; conversely, depletion of an shRNA indicated that gene ablation is detrimental for tumour cells and the target gene therefore may promote metastasis. To determine *in vivo-*specific effects, shRNA representation in cells grown in tissue culture (*in vitro*) was analysed at the same time points. We identified 27 genes where three different shRNAs targeted against the same gene were depleted by log_2_ fold change of 1 or greater under *in vitro* conditions (Figure S1B). A number of these genes have previously been shown to play an important role in cell survival, including *Eif5b, Mdn1, Rbm8a, Rrm1, Snrpd1*, and *Timeless* (Wang et al., 2015a). These shRNAs were also depleted in lung samples to a similar degree indicating a shared requirement for these genes under *in vitro* and *in vivo* conditions. We next sought to investigate lung-specific (*in vivo*-only) effects (Figure 1B, Supplementary Table 2) and selected a number of targets for further investigation. Notably, shRNAs against *Kras* were more depleted in cells grown *in vivo* than *in vitro* (Figure 1C), indicating the requirement for *Kras* is amplified in the secondary site microenvironment. Selected candidates were further validated in *in vivo* experiments, including *Pacs2, Mll3, Msi2*, and *Zdhhc20*. The efficiency of shRNA knock-down of these genes was assessed by quantitative PCR (Figure S1C-F), and by western blot for ZDHHC20 (Figure S1G). In functional assays, cells expressing shRNA targeting *Msi2* proliferated at the same rate in 2D culture as the control (Figure 1D), but their tumourigenicity *in vivo* was significantly reduced (Figures 1E and 1F). Similar results were observed with shRNA targeting *Mll3* (Figures 1G-I), *Pacs2* (Figures 1J-L), and *Zdhhc20* (Figures 1M-O), where the effect on tumour formation in the lung was several times higher than the effect in culture, suggesting a potential pro-metastatic role for these genes.

### ZDHHC20 validated as a novel mediator of PDAC metastasis

We chose to focus our further studies on ZDHHC20, a member of the ZDHHC family of *S*-acyltransferase enzymes, which is widely conserved between mouse and human (Salaun et al., 2020). ZDHHCs regulate membrane association and trafficking of a wide range of protein substrates in mammalian cells (Blanc et al., 2019) through long-chain fatty acid cysteine acylation. We undertook a CRISPR/Cas9 *Zdhhc20* gene knock-out (KO), and did not observe significant compensatory expression of other *Zdhhc* genes (Figures S2A and S2B). *Zdhhc20* KO (Figure 2A) did not affect cell proliferation in culture (Figure 2B), anchorage-independent growth (Figure S2C), colony formation (Figure S2D), or migration (Figure S2E). Strikingly, *Zdhhc20* KO almost completely inhibited secondary site outgrowth of PDAC cells in the lung (Figures 2C, 2D, S2F) and liver (Figures 2E and 2F). Long-term observation of tumour growth in mice injected with KO cells identified rare macroscopic tumours at a later time-point (Figure S2G). This suggested an adaptation mechanism or a very low seeding-density of metastatic cells due to the initial attrition by KO of *Zdhhc20*. We also generated KO cells in another PDAC-derived cell line (FC1199) (Figure 2G), and observed a similar phenotype *in vitro* (Figures 2H, S2H-S2J), and a loss of tumourigenicity *in vivo* (Figure 2I). To investigate the relevance of *Zdhhc20* in human PDAC, we compared overall survival probability as a function *Zdhhc20* gene expression. Analysis of TCGA datasets revealed that high *Zdhhc20* expression is associated with significantly worse survival in PDAC patients (Figure 2J). Additionally, *Zdhhc20* was identified as a part of a prognostic gene expression classifier in patients with resectable disease, being upregulated in short-term survivors compared with long-term survivors (Birnbaum et al., 2017).

**Figure 2.**
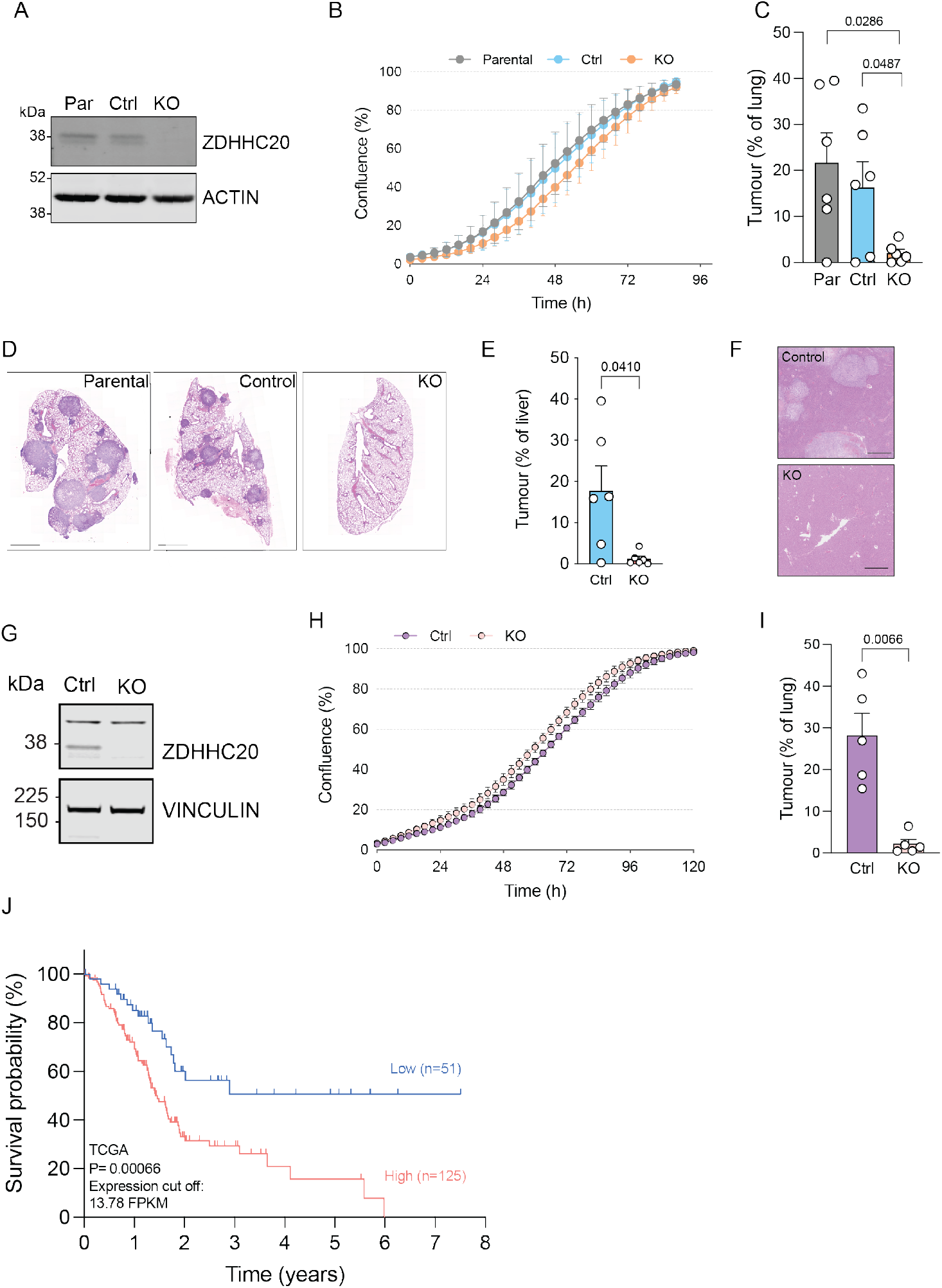
Validation of ZDHHC20 as mediator of PDAC metastasis *in vivo*. **(A)** Western blot image depicting ZDHHC20 level in parental PDA530Met cells, control cells, and CRISPR KO clone of Zdhhc20. **(B)** Proliferation curves of parental, control, and *Zdhhc20* KO in adherent 2D culture (n=3). **(C)** Quantification of tumour area in the lungs of mice injected with parental, control, and *Zdhhc20* KO cells. **(D)** H&E staining of mouse lung tissue at day 28 after i.v. injection of parental, control or CRISPR *Zdhhc20* KO cells. Scale bar, 1000 µm. **(E-F)** Quantification of tumour area **(E)** and H&E staining of liver tissue **(F)** following intrasplenic injection of control or *Zdhhc20* KO cells. Scale bar, 1000 µm. **(G)** Western blot image of ZDHHC20 in control and KO cells generated by Cas9/CRISPR in FC1199 mouse PDAC cell line. **(H)** Growth curves of control and FC1199 *Zdhhc20* KO cells *in vitro* (n=3). **(I)** Quantification of tumour area in the lungs following i.v. injection of FC1199 control and *Zdhhc20* KO cells. **(J)** Kaplan-Meier survival curve of PDAC patients from the TCGA dataset separated according to best cut-off *ZDHHC20* expression. Data are shown as mean + S.E.M. An unpaired t-test with Welch’s correction (does not assume equal standard deviations) was used in **(C), (E)**, and **(I)**. P-values are shown above the comparison lines.

### Tumourigenicity of *Zdhhc20* KO cells is dependent on NK cells

As *Zdhhc20* KO cells did not form tumours when injected intravenously into immunocompetent mice, we sought to investigate whether certain components of the immune system could mediate this phenotype. When injected into Rag1^-/-^ mice, which lack B and T cells, we observed a replication of the phenotype seen in wild-type animals (Figure 3A). In contrast, the KO cells readily formed tumours in more immunocompromised animals, which lack B cells, T cells, and natural killer (NK) cells (Figures 3B and 3C), demonstrating that the lack of metastatic ability is not a cell-autonomous feature of *Zdhhc20* KO cells, but is dependent on the presence of a functional immune system, primarily NK cells. To investigate this observation further, we carried out NK cell depletion using neutralising antibodies in combination with the injection of tumour cells (Figures 3D, S3A, and S3B). Wild-type mice (129S background) are fully immunocompetent, and depletion of NK cells in this model was achieved using anti-asialo GM1 antibody because of the lack of anti-Nk1.1 reactivity in this strain (Carlyle et al., 2006). Indeed, depletion of NK cells using neutralising antibodies rescued the ability of KO cells to form tumours in the lung (Figures 3E). The phenotype rescue was not complete, potentially due to some residual NK cell activity in the depletion setting or another component of innate immunity also contributing to targeting of *Zdhhc20* KO cells.

**Figure 3.**
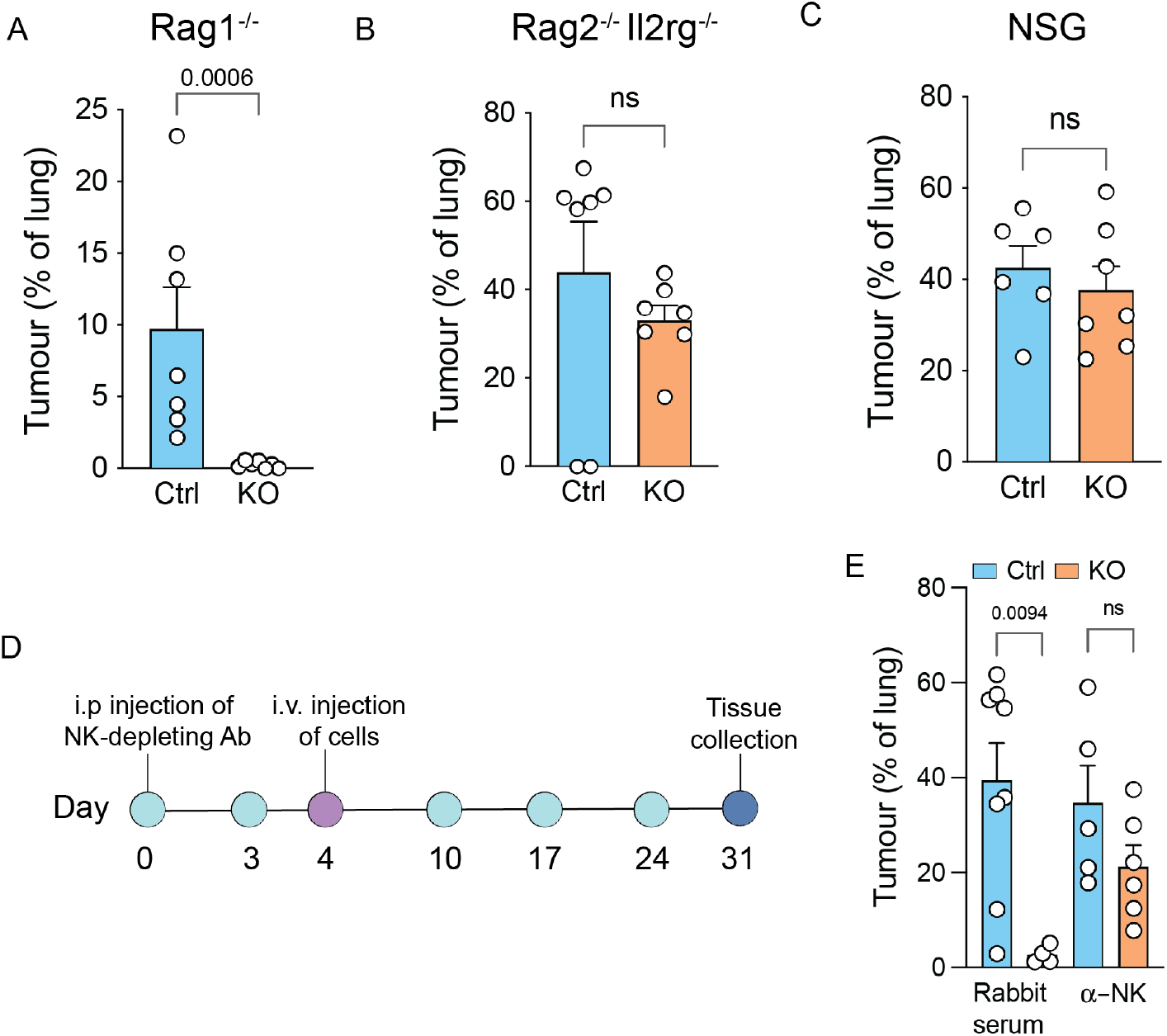
Tumourigenicity of *Zdhhc20* KO cells is dependent on NK cells. **(A-C)** Quantification of tumour area in the lungs of mice injected with control cells and *Zdhhc20* KO cells in Rag1^-/-^ **(A)**, Rag2^-/-^Il2rg^-/-^ **(B)**, and NOD scid gamma **(C)** backgrounds. **(D)** Schematic of the NK cell depletion protocol. **(E)** Analysis of tumour area in wild type 129S mice treated with rabbit serum or NK cell-depleting serum (anti-asialo GM1), injected with control or *Zdhhc20* KO cells. Mann-Whitney test was used for comparisons in **(A), (B), (C)**, and **(E)**.

### *Zdhhc20* KO inhibits metastasis in a murine model of PDAC

We sought to complement findings from the experimental metastasis setting by investigating the effect of ZDHHC20 loss in a genetically engineered mouse model. We generated a novel mouse line in which the exons coding for the active site of the ZDHHC20 enzyme (exons 5-7) are flanked by loxP sites (Figure S4A). This region is excised by Cre recombinase, resulting in nonsense-mediated decay of the transcript, while any potentially translated protein would be truncated and catalytically inactive. The constitutive whole-body KO was generated by crossing *Zdhhc20*^fl/fl^ with a *PGK-Cre* line (Lallemand et al., 1998), and KO offspring were viable and showed no adverse phenotype or histological abnormalities as adults (Supplementary Table 3). Recombination efficiency and specificity in the pancreas were validated after crossing with the line expressing a pancreas-specific *Pdx1-Cre* transgene (Figure S4B).

We then introduced *Zdhhc20*^fl/fl^ allele (Z) into the classical KPC (*LSL-Kras*^*G12D/+*^; *LSL-Trp53*^*R172H/+*^; *Pdx-1-Cre)* model (Figure S4C), characterised by pancreatic adenocarcinoma and distant metastasis to the liver and lung, which resembles human disease (Hingorani et al., 2005). In this model, pancreatic tumourigenesis is initiated by the endogenous expression of mutant *Kras* (G12D) in the developing pancreas, driven by the expression of the *Pdx1-Cre* transgene. In combination with the expression of mutant *Trp53* (R172H) and loss of the wild-type allele, these animals develop invasive metastatic disease. We first assessed the overall survival of tumour-bearing animals with a pancreas-specific deletion of *Zdhhc20*. No change in survival among wild type (WT), heterozygous (HET, one floxed *Zdhhc20* allele), and homozygous (HOM, both *Zdhhc20* alleles floxed) KPCZ mice was observed (Figure 4A). Survival in this model is mainly affected by the growth of the primary tumour, suggesting that *Zdhhc20* expression is not essential for primary PDAC growth. Similarly, there was no difference in the overall histology of these tumours in different *Zdhhc20* genotypes (Figures 4B, S4D).

**Figure 4.**
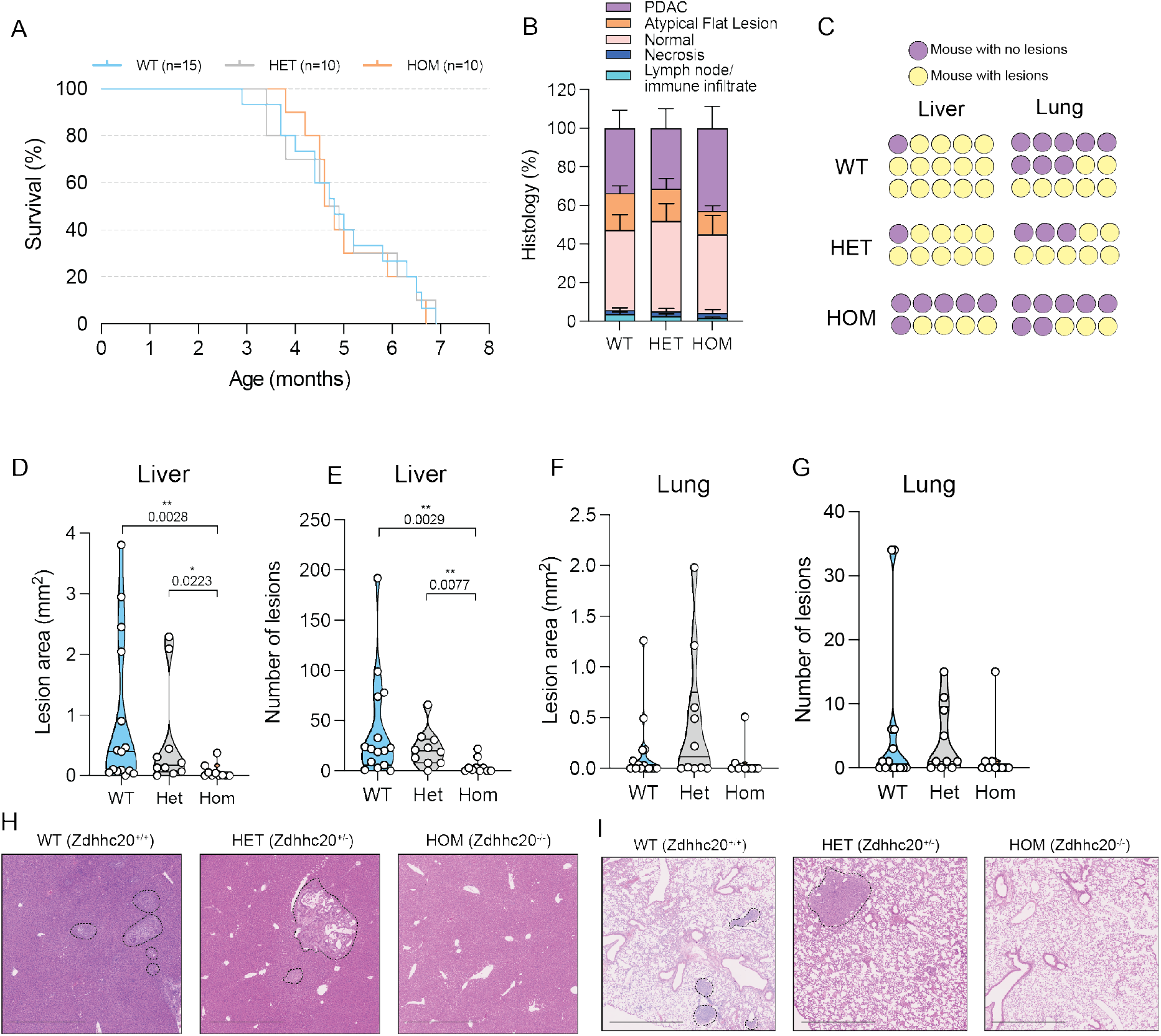
*Zdhhc20* KO inhibits metastasis in KPC mouse model of pancreatic cancer. **(A)** Survival curve of wild type KPC (WT, *Zdhhc20*^+/+^, 6 males, 9 females), heterozygous (HET, pancreas-specific *Zdhhc20*^+/-^, 6 males, 4 females), or homozygous (HOM, pancreas-specific *Zdhhc20*^-/-^, 4 males, 6 females) mice. **(B)** Primary tumour histology in KPCZ mice. **(C)** Frequency of metastatic lesions in liver and lung tissue of KPCZ mice, based on H&E staining analysis. **(D-E)** Quantification of liver lesion area **(D)** and number **(E)** in KPCZ mice. **(F-G)** Quantification of lung lesion area **(F)** and number **(G)** in KPCZ mice. **(H-I)** Representative images of liver **(H)** and lung **(I)** H&E sections from KPCZ mice, Scale bar, 1000 µm. Mann-Whitney test was used for comparisons in **(D)** and **(E)**.

We then assessed H&E section of liver and lung from these mice and quantified microscopic and macroscopic lesions in these tissues. As previously reported for a murine model of PDAC, the metastatic burden was highly-variable (Maddipati et al., 2022). Despite this inherent variability, a lower frequency of liver lesions was observed in *Zdhhc20* HOM compared with HET or WT mice, with metastases found in 40%, 90%, and 93% of mice, respectively (Figure 4C). A similar trend was observed for lung lesions, although with a reduced incidence of lesions, given that lungs are a site with lower frequency of metastatic spread of PDAC compared with the liver. Quantification of total liver lesion area and number showed a profound reduction in HOM animals, with a similar, but non-significant trend observed in the lung (Figures 4D-4G). Histology of these lesions varied from differentiated to invasive, highly-infiltrated with stromal response (Figures 4H, 4I), similar to previous reports (Zhong et al., 2017).

To validate the pancreatic tissue origin of these lesions, we generated a KPC line with a fluorescent LSL-tdTomato reporter in the *Rosa26* locus, which enabled tracing of metastatic cells and lesions (Figure S4E). Together, these results demonstrate the importance of *Zdhhc20* expression for the metastatic spread of PDAC, but not for primary tumour growth.

### Proteomic identification of ZDHHC20 effectors in metastatic PDAC cells

We next focussed on delineating the molecular mechanism underlying the requirement of ZDHHC20 for metastasis of pancreatic cancer. ZDHHC20 is an integral membrane protein that catalyses protein *S*-acylation, often termed palmitoylation, a post-translational modification in which a long-chain fatty acid such as palmitate (C16:0) is transferred to the target protein (Stix et al., 2020). In humans, ZDHHC20 is a part of a family of 23 *S*-acyltransferases, recognised by their characteristic zinc finger Asp-His-His-Cys (ZDHHC) motif (Korycka et al., 2012). Direct determination of the substrates of a given ZDHHC by standard proteomic methods is confounded by the great diversity and complexity of the *S*-acylated proteome, numbering >3000 S-acylation sites in mammalian cells. We therefore used a novel chemical genetic approach to map the protein substrates of ZDHHC20 (Ocasio, Baggelaar, Sipthorp et al., unpublished). This approach exploits an engineered ZDHHC20^Y181G^ mutant and a corresponding click (alkyne) tagged ‘bumped’ fatty acid probe that is selectively loaded on the mutant ZDHHC and then transferred to the substrates of the enzyme in cells (Figures 5A and S5A). WT ZDHHC20 and Y181G mutant were stably overexpressed in PDA530Met cells (Figure 5B) and selective loading of the chemical probe on the Y181G mutant was confirmed (Figure S5B). Substrate profiling through click chemistry ligation to a biotin label and affinity purification followed by quantitative proteomic analysis by nanoflow chromatography-MS/MS identified a total of 124 potential protein substrates of ZDHHC20 (Figure 5C, Supplementary Table 4). To generate a complementary view of changes in protein palmitoylation upon ZDHHC20 loss, we also carried out an analysis of the palmitoylated proteome by metabolic labelling with 17-ODYA and click chemistry-enabled proteomic analysis following knock-down (KD) of *Zdhhc20*, identifying 58 differentially represented palmitoylated proteins in *Zdhhc20* KD cells. We compared this list with ZDHHC20 substrates (Figure S5C, Supplementary Table 5), revealing a change in the *S*-acylated proteome beyond a direct impact on substrates. Loss of ZDHHC20 activity has the potential to profoundly influence protein-protein interactions at the membrane, as well as *S*-acylation between ZDHHCs themselves (Zmuda and Chamberlain, 2020), with the potential to strongly influence presentation of cell surface receptors and proteins beyond the direct substrates of ZDHHC20. With ZDHHC20 predominantly expressed in vesicles and the plasma membrane (Human Protein Atlas; Thul et al., 2017), we specifically explored the changes in cell surface proteome using a cell surface biotinylation assay (Scheurer et al., 2005) coupled with mass spectrometry proteomics and identified differentially represented proteins between parental and KO cells (Figures 5D and 5E, Supplementary Table 6).

**Figure 5.**
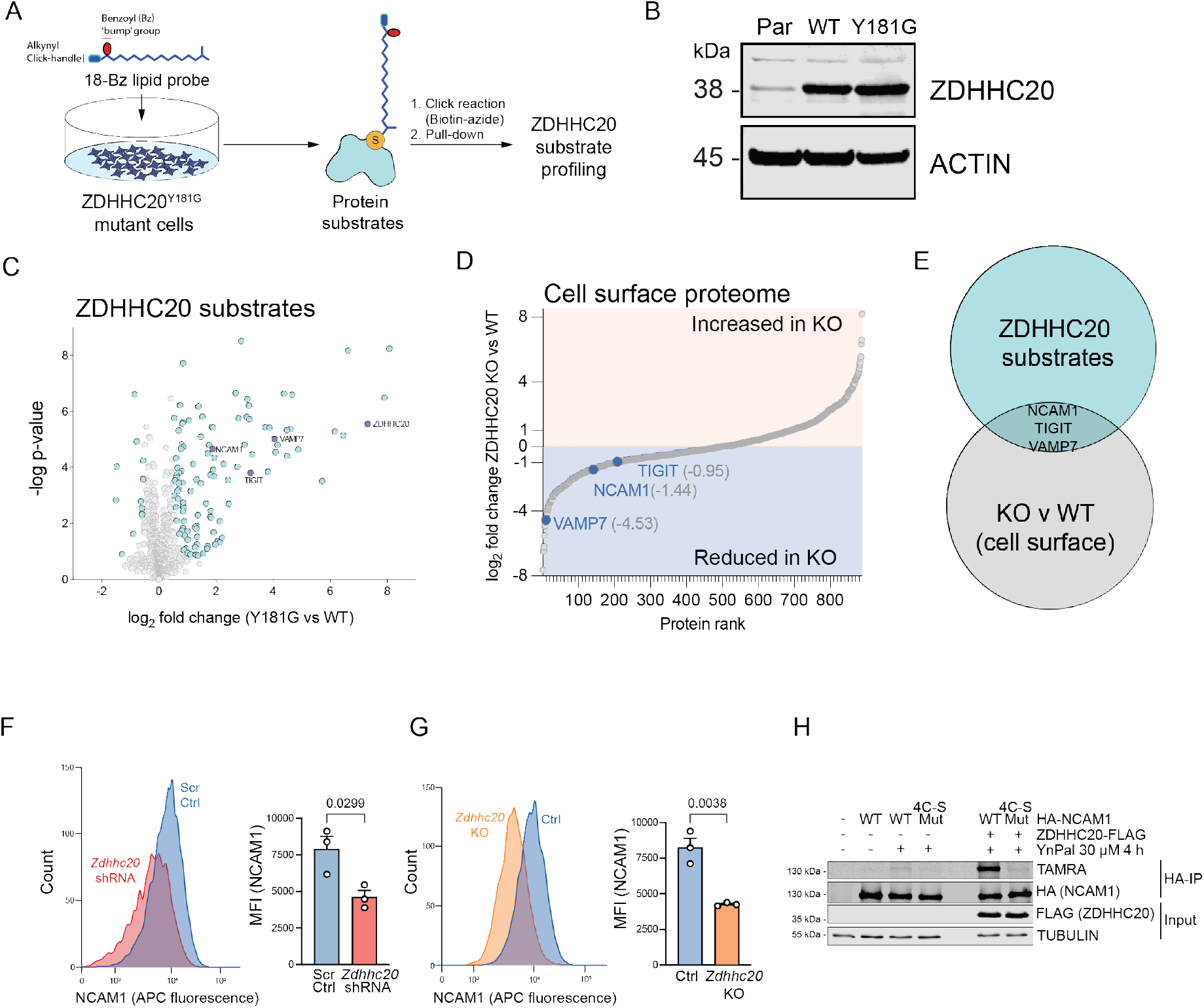
Identification of ZDHHC20 substrates. **(A)** Schematic of ZDHHC20 substrate profiling protocol. **(B)** Western blot of ZDHHC20 in parental, ZDHHC20^WT^-overexpressing and ZDHHC20^Y181G^-overexpressing cells. **(C)** Enrichment of palmitoylated proteins in Zdhhc20^Y181G^-overexpressing versus ZDHHC20^WT^-overexpressing cells following ‘bumped’ alkyne palmitate probe treatment and palmitoylated proteome pulldown (n=4, false discovery rate (FDR) = 5%). **(D)** S-plot of differentially-represented cell surface proteins in *Zdhhc20* KO v WT (Par). **(E)** Overlap of ZDHHC20 substrates with the differentially-represented cell-surface proteins in KO v WT cells. **(F)** FACS histogram and quantification of NCAM1 median fluorescence intensity (MFI) in Scr and *Zdhhc20* shRNA cells. **(G)** FACS histogram and quantification of NCAM1 median fluorescence intensity (MFI) in control and *Zdhhc20* KO cells. **(H)** Palmitoylation assay of WT NCAM1 (WT) and 4C-S NCAM1 (4C-S Mut). Data are shown as mean + S.E.M. Two-tailed t-test was used for data in **(F)** and **(G)**. P-values are shown for each comparison.

Overall, we identified three novel substrates of ZDHHC20 with decreased cell surface level: Neural Cell Adhesion Molecule 1 (NCAM1), Vesicle-associated membrane protein 7 (VAMP7) and T cell immunoreceptor with Ig and ITIM domains (TIGIT). The downregulation of these proteins at the cell surface was not due to transcriptional regulation, as the level of mRNA transcript remained unchanged in KO cells (Figure S5D). Despite the downregulation of VAMP7 at the cell surface, the size and the number of extracellular vesicles was not affected (Figure S5E). We validated the decrease of NCAM1 in both shRNA and KO cells by flow cytometry (Figures 5F and 5G). In biochemical assays, 4C-S mutant NCAM1, where the four cysteine residues in the cytoplasmic tail of the protein are mutated into serine, cannot be *S*-acylated by ZDHHC20 (Figures 5H and S5F).

## Discussion

We identified several PDAC-associated genes which mediate metastasis *in vivo* by comparing the representation of shRNA in cells grown in culture and in lung tumours following intravenous injection into immunocompetent animals. This design helps to identify novel genes relevant for the metastatic process, particularly in the context of interaction with components of the immune system. Here, we discovered a protein *S*-acyltransferase, ZDHHC20, as one such mediator. Using mouse PDAC cell lines PDA530Met and FC1199, we show that metastatic potential is significantly reduced upon the loss of ZDHHC20, without an apparent effect on the phenotype of cells grown in culture. In a mouse model of pancreatic cancer, we show that despite having no effect on primary tumour, a complete loss of ZDHHC20 inhibited metastasis to the liver and lung, supporting the findings from the initial shRNA screen.

Interestingly, we found that *Zdhhc20* KO cells readily form tumours in immunocompromised mice. This effect is at least in part mediated by NK cells, as NK cell depletion in wild-type mice promoted tumourigenicity of KO cells relative to control cells. We hypothesised that this dependency on NK cells might be due to the alterations of the cell surface proteome of KO cells, given that ZDHHC20 is predominantly expressed in the vesicles and the plasma membrane. By combining a new technology of ZDHHC20 chemical genetic substrate profiling with cell surface proteomics, we propose several candidates which could mediate this phenotype, including through interaction with NK cells and other components of the immune system. In particular, identification of novel protein substrates of this enzyme in metastatic mouse PDAC cells expands the map of pathways that may be regulated by *S*-acylation (also often termed “palmitoylation”), and which might be therapeutically targeted in future to control PDAC metastasis.

The role of palmitoylation in regulating cancer-associated phenotypes is an emerging area of research, where multiple ZDHHC *S*-acyltransferases have been shown to regulate processes including cell proliferation, polarity, transcriptional activation and invasion (Ko and Dixon, 2018; Zhang et al., 2020). Previous studies suggest that ZDHHC20 may play a role in cellular transformation and cancer (Draper and Smith, 2010). Studies in lung and breast cancer cells have found that epidermal growth factor receptor (EGFR) may be palmitoylated by ZDHHC20 (Runkle et al., 2016), and that blocking EGFR palmitoylation by ZDHHC20 inhibition may supress mutant KRAS-induced primary lung tumour formation (Kharbanda et al., 2020). However, in melanoma, this enzyme has been suggested to *S*-acylate melanoma cell adhesion molecule (MCAM), leading to increased cell invasion (Wang et al., 2015b), and different tissue type effects may be due to differing ZDHHC20 substrate profiles of different cell lineages.

We suggest that, since constitutive *Zdhhc20* KO mice are viable and of normal tissue histology, targeting this enzyme or its substrates presents a novel potential therapeutic approach to limit or prevent metastatic spread in PDAC.

## Methods

### Cell lines

Mouse metastatic pancreatic ductal adenocarcinoma PDA530Met cell line (kindly provided by David Tuveson, Cold Spring Harbor Laboratory) was grown in Dulbecco’s Modified Eagle Medium (DMEM) with L-glutamine, 10% foetal bovine serum (FBS) and antibiotics (Penicillin/Streptomycin, 100 U/mL). PDA530Met cells were derived from a liver metastasis of a male mouse of the *Kras*^*G12D/+*^; *Trp53*^*R172H/+*^; *Pdx1-Cre* (KPC) model of pancreatic cancer (Hingorani et al., 2005), of a mixed 129/SvJae and C57Bl/6 background. FC1199 cell line was derived from a primary pancreatic tumour of a male KPC mouse of an inbred C57Bl/6 background (kindly provided by David Tuveson, Cold Spring Harbor Laboratory).

### Mice

Mice used for intravenous injection experiments were 8-12 weeks old males, sourced internally at the Francis Crick Institute or from a commercial supplier (Charles River). The mice were housed in a specific-pathogen-free (SPF) facility, under controlled conditions with a 12 h light/dark cycle. The strains used in experiments were 129S6/SvEv, Rag1 KO (Rag1tm1Mom), Rag2^-/-^ Il2rg^-/-^, and NOD scid gamma. All animal experiments were done in compliance with the regulations of the UK Home Office and approved by the Animal Welfare and Ethical Review Body at the Francis Crick Institute or approved by the Institutional Animal Care and Use Committee (IACUC) at Cold Spring Harbor Laboratory.

### *Zdhhc20* KO mouse line

A conditional *Zdhhc20* KO mouse model was generated on a C57Bl/6 background by a commercial service provider (Biocytogen). The exons 5-7 of mouse *Zdhhc20* (NCBI Gene ID 75965, coding the catalytic site of the enzyme) were flanked by loxP sites inserted into the non-conserved regions of introns 4 and 7. The targeting construct containing the loxP sites and homology arms was inserted using CRISPR/EGE™ (CRISPR/Extreme Genome Editing) technology. The 5’-sgRNA target sequence was TATGCTATATCACCAAGCATTGG, and the 3’-sgRNA target sequence was CGGTAACCGGGAGTGACCTCTGG. Following F0 generation screening and genotyping, F1 mice were validated for germline transmission, rederived at the Francis Crick Institute, and bred to homozygosity for the *Zdhhc20*^fl/fl^ allele. Validation of targeting and sequence were confirmed in-house. To test the phenotype of a constitutive, whole-body deletion of *Zdhhc20*, the mice were crossed with the PGK-Cre line (Tg(Pgk1-cre)1Lni, MGI:2178050).

The *Zdhhc20*^fl/fl^ mice were then crossed with *Pdx1-Cre* line (Tg(Pdx1-cre)6Tuv, MGI:3032531), *LSL-Kras* G12D line (tm4Tyj, MGI:2429948), and *LSL-p53* R172H line (tm2tyj, MGI:3039263), all previously backcrossed to C57Bl/6 background. The offspring of these crosses were bred to generate KPCZ (*LSL-Kras*^*G12D/+*^; *LSL-Trp53*^*R172H/+*^; *Pdx1-Cre, Zdhhc20*^*fl/fl*^) mice used for pancreatic tumourigenesis studies. For experiments involving tracing tumour cells using a fluorescent reporter, tdTomato, the KPCZ mice were further crossed with *Rosa26-CAG-LSL-tdTomato* line (tm14(CAG-tdTomato)Hze, MGI:3809524), to generate KPCZT mice. Genetic monitoring of inbreeding status was carried out using MiniMUGA background analysis (Transnetyx), and the mice confirmed as being close to inbred. Diagnostic SNPs indicated the presence of the background strain groups C57BL/6 and the substrains B6N-Tyr<c-Brd>/BrdCrCrl, C57BL/6J.

### Lentivirus production

HEK293T cells were lipofected with the pLKO-puro plasmid, pCMV delta R8.2, and pCMV-VSV-G plasmid, followed by the change of medium after 8 hours. On day 4 post-infection, the supernatant was collected and passed through a 0.45 µm filter.

### *In vivo* shRNA screen

An shRNA library targeting 284 pancreatic cancer related genes was generated using the pLKO vector backbone (Sigma). Each gene was targeted with 3-4 shRNA producing a library size of 1013 shRNA. Nineteen shRNA pools were generated and each contained 8 control shRNA to assess consistency across pools. Virus for each pool was generated and used to infect target PDA530Met cells at an MOI of 0.3. Cells were selected for 4 days with puromycin (3 µg/ml) and then 5 × 10^5^ cells were injected via tail vein into mice (3 mice per pool). A cell pellet was also generated to provide a Day 0 sample of each pool. After 28 days, lung samples were harvested and all tumours within the same mouse were pooled to generate a pooled lung DNA sample. In parallel, cells were cultured *in vitro* in triplicate over the same time-frame. An shRNA representation of 2000X was maintained at each step throughout the screen.

### DNA isolation and sequencing

Genomic DNA extraction of Day 0, *in vitro* and *in vivo* samples was carried out using Gentra Puregene Tissue Kit (Qiagen). A two-step PCR was carried out to amplify shRNA sequences from the genomic DNA samples. Sequencing adapters and multiplexing barcodes were incorporated into the PCR primer (Molina-Arcas et al., 2019). PCR products were purified using MinElute PCR Purification Kit (Qiagen), and DNA concentration was determined using Bioanalyser (Agilent). Next generation sequencing was carried out to determine the representation of each shRNA in the samples.

### RNA isolation and sequencing

Total RNA was isolated using GenElute™ Mammalian Total RNA Miniprep Kit (Sigma-Aldrich) from approximately 5 million cells per biological sample of PDA530Met cells, control and KO clones (n=3). Sample quality of isolated RNA was assessed using 2100 Bioanalyzer (Agilent). The libraries were prepared using KAPA mRNA HyperPrep kit (Roche), and sequenced on HiSeq 4000 system (Illumina) at 75 bp single read (25M reads/sample).

### Western blotting

Protein extracts were prepared in a lysis buffer (1% Triton X-100, 0.1% SDS in PBS), containing protease and phosphatase inhibitors (Sigma-Aldrich). Following cell lysis, the extracts were incubated on ice for 20 minutes, with gentle agitation. The samples were centrifuged at 10,000xg and supernatants collected for downstream analysis. Protein concentration was quantified using DC protein assay (Bio-Rad) according to manufacturer’s instructions. The membrane was blocked with Odyssey blocking buffer (LI-COR Biosciences) or 5% fish gelatine in TBS for 1 hour at RT, followed by an overnight incubation with the primary antibodies diluted in Odyssey buffer containing 0.2% Tween-20. Secondary antibodies (680RD donkey anti-mouse and 800CW donkey anti-rabbit, 1:20,000, LI-COR Biosciences) were diluted in Odyssey buffer containing 0.1% Tween-20 and 0.01% SDS, and incubated for 1 hour at RT in the dark. The membranes were imaged using Odyssey CLx Infrared Imaging System and the images processed with Image Studio Lite (LI-COR Biosciences).

### ShRNA knockdown of *Msi2, Mll3, Pacs2*, and *Zdhhc20*

PDA530Met cells were transduced with control Scr shRNA (Non-Mammalian shRNA Control, SHC002, Sigma-Aldrich) or two independent shRNAs directed against the target gene: *Msi2* (TRCN0000071976 and TRCN0000071974), *Mll3* (TRCN0000238934 and TRCN0000238936), *Pacs2* (TRCN0000250359 and TRCN0000250357), *Zdhhc20* (TRCN0000328625 and TRCN0000328698, Sigma-Aldrich). Following selection with puromycin (3 µg/mL) for 4 days, 1 × 10^5^ cells were injected via tail vein into mice.

### Identification of *Zdhhc20* knock-down cell palmitome

Parental PDA530Met cells were seeded at a density of 30 000 cells per 10 cm plate. After 24 hours, fresh medium with polybrene (10 µg/mL) was added, and cells were infected with lentivirus expressing Scr shRNA or two independent shRNA targeting *Zdhhc20*. Selection was carried out using 3 µg/mL puromycin, and cells were grown in SILAC heavy media, SILAC medium media or normal media. After 7 days of selection and growth in SILAC-media, cells were treated overnight with a palmitic acid analogue (50 µM) to label newly palmitoylated proteins. The samples were then lysed in cell lysis buffer (1% Triton X-100, 0.1% SDS in PBS, containing protease and phosphatase inhibitors) and protein concentration was determined via DC assay. Lysates were pooled and a Click-IT assay was carried out to facilitate binding of Biotin azide with 17-ODYA incorporated proteins. Proteins were precipitated using methanol/chloroform, resuspended, and biotin-conjugated proteins were pulled-down using NeutrAvidin beads (Thermo). Mass spectrometry analysis was carried out to identify palmitoylated proteins. Raw files were processed in MaxQuant software and H/L and H/M from Protein Groups table imported into Perseus software (basic filtering and transformation, Significance B).

### CRISPR-mediated *Zdhhc20* gene knock-out

Guide RNAs targeting exon1 of *Zdhhc20* were pre-designed and ordered in a pU6 vector backbone (Sigma Aldrich). Two gRNA plasmids and pCas9 (D10)-GFP plasmid were nucleofected into cells using Basic Nucleofector Kit for Primary Mammalian Epithelial Cells (Lonza) according to the manufacturer’s instructions. The nucleofected cells were allowed to recover for 24-48 h, followed by Fluorescence Activated Cell Sorting (FACS) of GFP^+^ single cells using DAPI or Hoechst 33258 as a viability marker. The knock-out clones were validated by Western blot, and by using PCR followed by Sanger sequencing.

### Flow cytometry

For sorting, cells were detached using 0.25% trypsin (Sigma), resuspended in DMEM with 10% FBS, spun down for 3 minutes at 1200 rpm at a benchtop centrifuge, resuspended in PBS, and filtered through a 40 µm cell strainer before sorting. Live single-cells were sorted directly into medium-filled wells of a 96-well plate using either BD FACSAria™ Fusion Flow Cytometer (BD Biosciences) or MoFlo XDP Cell Sorter (Beckman Coulter).

For all other experiments, cells were analysed using BD LSRFortessa™ (BD Biosciences). For cell line flow cytometry analysis, the cells were detached using StemPro™ Accutase™ Cell Dissociation Reagent (Thermo Fisher), and further processed in the same manner as for sorting. For NK cell analysis, the spleen was first pushed through a 70 µm cell strainer, rinsed with PBS, and centrifuged for 5 min at 4ºC. The pellet was resuspended in ACK lysing buffer (Thermo Fisher), to lyse erythrocytes. After the washing steps, the cells were incubated with anti-CD16/CD32 antibody (1:100, BD Biosciences) for 10 min on ice, followed with the addition of anti-CD45 antibody (1:600, PerCP, Biolegend) and Nkp46 antibody (1:40, PE, Biolegend) for 30 min on ice. The cells were washed again, and DAPI (1 µg/mL final concentration) added before analysis. The washing steps and final resuspension were carried out with a buffer solution containing 2 mM EDTA and 0.5% BSA in PBS (pH 7.2).

### Cloning of *Zdhhc20* constructs

The *Zdhhc20*^*WT*^-attb plasmid was generated using InFusion cloning. Briefly, the attb-containing part of pX28 plasmid backbone (#46850, Addgene) was PCR amplified, and joined with individually amplified PGK promoter, *Zdhhc20* CDS, BGH pA sequence, SV40 Promoter-Blasticidin-S deaminase sequence, and SV40 pA sequence in a single InFusion cloning reaction. The correct assembly of the plasmid was confirmed by Sanger sequencing. The construct containing Y181G-coding point mutation was generated by inverse PCR of *Zdhhc20*^*WT*^ plasmid using CloneAmp HiFi PCR Premix (Takara) and joining the ends of the PCR product using InFusion cloning.

### Cloning of Ncam1 constructs

A pCMV3 plasmid containing cDNA for mouse neural cell adhesion molecule 1 (Ncam1, transcript variant 3, NCAM1-180) was used for cloning and expression (MG56977-UT, SinoBiological). The cysteine sites in the cytoplasmic region (SwissPalm, entry P13595) were selected for mutation into serine: Cys734, Cys740, Cys745, and Cys751. The 4C-S Ncam1 mutant plasmid was generated by joining a synthesised oligonucleotide fragment containing all four mutations with the plasmid backbone using InFusion enzyme mix. The hemagglutinin (HA) tag was added at the N-terminus of WT and 4C-S NCAM1 by joining an HA tag-coding oligonucleotide fragment with the backbone of either WT or mutant Ncam1 plasmid.

### Generation of *Zdhhc20*-overexpressing cells

The PhiC31 integrase was utilised to generate a cell line overexpressing *Zdhhc20* at a near-endogenous level. The plasmid expressing the PhiC31 integrase was transfected using Lipofectamine 3000 (Life Technologies), in combination with the *Zdhhc20* plasmid containing a PGK promoter, attb recombination sequence, and the Blasticidin resistance cassette. The ratio of PhiC31 and *Zdhhc20*-attb plasmid was 50:1 (w/w). Following lipofection, the cells were plated at a low density in a 5 µg/mL Blasticidin-containing medium. Single-cell colonies were picked using colony cylinders and expanded. Integration of the *Zdhhc20* cassette was checked by PCR and overexpression confirmed by Western Blot.

### Cell proliferation assays

To analyse proliferation, shRNA knockdown cells were plated at 2 × 10^3^ cells per well in a 24-well plate. After 6 days cells were stained with crystal violet stain (0.5% crystal violet in 2% ethanol solution) for 30 mins. The crystal violet dye was dissolved in 10% acetic acid solution (1 mL per well), and absorbance at 595 nm was measured using a plate reader (Tecan). Proliferation of all other cell lines was determined using an IncuCyte system (Essen BioScience), where 1 × 10^4^ cells were plated in a 24-well plate and confluence measured automatically every 3 hours, for up to 5 days.

### Anchorage-independent growth assay

Cells were trypsinised, passed through a 40 µm filter, and plated in a low attachment 96-well plate (Corning) at a concentration of 1 × 10^4^ cells/well. Following a 72 h incubation, viability of the cells was analysed using CellTiter-Glo 3D assay (Promega), according to the manufacturer’s instructions.

### Colony formation assay

This experiment was carried out as previously described, as a 3D on-top assay (Lee et al., 2007). A total of 2500 single cells were plated on a layer of Matrigel (Corning) in a 48-well plate, and covered with a 10% Matrigel in cell culture medium. The colonies were allowed to form over 4 days, with 10% Matrigel medium being replenished after 2 days. The number of colonies was analysed using an automated colony counter (Oxford Optronics).

### Cell migration assay

To assess cell migration in 2D, a scratch-wound assay was utilised. The cells were seeded in an IncuCyte ImageLock 96-well plate at a cell density of 2 × 10^4^ cells/well and incubated for 24 h to reach full confluency. The scratch was created using the WoundMaker (Essen BioScience), followed by a wash to remove cell debris. The wound closure was monitored using the IncuCyte system.

### Intravenous injections

For shRNA screen experiment, PDA530Met cells were transduced with control Scr shRNA or two independent shRNA directed against the target gene. Following selection with puromycin (3 µg/mL) for 4 days, 1 × 10^5^ cells were injected via tail vein into mice and lungs collected after 4 weeks. For validation experiments using CRISPR KO clones, a total of 7 × 10^4^ cells PDA530Met cells or 2 × 10^5^ FC1199 cells were injected, and lungs collected after 3-4 weeks.

### Splenic injections

For splenic injections, the cells were prepared in PBS (5 × 10^5^ cells in 50 µL). The mice were anaesthetised and the spleen was divided in two parts using a cautery pen, cells injected into one part, allowed to rest for a few seconds, after which that part of spleen was removed. The livers were collected after 3 weeks.

### Histology

Tissues were fixed with 10% neutral buffered formalin solution for 24 hours, transferred to 70% ethanol, paraffin embedded and stained with H&E stain. For shRNA screen validation experiments, tumour area and tumour number were measured using Nikon NIS-Elements microscope image analysis software. For CRISPR KO experiments, H&E slides were first scanned using Zeiss Axio Scan.Z1 slide scanner (Carl Zeiss), followed by an automated tumour/lung area analysis using StrataQuest image processing software (TissueGnostics).

### Survival curve of KPCZ mice

The humane endpoint criteria for KPCZ mice were one or more of the following symptoms: persistent hunched posture, piloerection, laboured breathing, sudden 20% weight loss of original body weight or 15% weight loss maintained for 48 hours, with or without the presence of palpable abdominal mass or ascites. Some animals developed tumours on the face, oral, and anogenital area. In these cases, the presence of tissue damage and bleeding in these areas was a contributing factor for culling. The animals where this was the only indicator for culling were excluded from the analysis.

### Histology analysis of KPCZ tissues

The H&E sections of pancreas, liver, and lung tissue were scanned using Zeiss Axio Scan.Z1 slide scanner (Carl Zeiss). The pancreas sections were assessed by two board certified veterinary pathologists (A.S-B. & S.L.P), and relevant histopathological categories annotated. The annotations were used to train pixel classifier in QuPath (version 0.3.2) in order to automatically quantify specific microscopic areas in the entire section and across multiple samples. The pixel classifier used was Random Trees (RTrees), at low/very low resolution, using haematoxylin and eosin channels. For samples where H&E intensity differed between batches, a classifier tailored to the sample was trained instead, to avoid misidentification. Background pixel class was subtracted and all other pixel classes added together to calculate the total area of the pancreas/tumour.

The lesions in the liver and lung were scored manually using the one of the following criteria: regions of glandular or cystic structures demarcated from the surrounding tissue, regions of high-cellularity and heterogeneity demarcated from the surrounding tissue. The H&E sections of livers and lungs from wild-type littermates were used as a benchmark for defining the normal histology of these tissues.

### Analysis of KPCZT mice

The pancreas, liver and lung of KPCZT mice were collected and fixed in freshly-made 4% formaldehyde in PBS for 36 hours, after which they were transferred into 15% sucrose for 24 hours, and 30% sucrose in PBS for a minimum of 48 hours, all at 4ºC. The tissue was then embedded in OCT and sectioned. Frozen sections were thawed at room temperature, mounting medium with DAPI added (Vector Laboratories), coverslipped, edges sealed with nail-varnish and imaged.

### ZDHHC20 substrate identification

PDA530Met cells with overexpressed ZDHHC20^WT^ or ZDHHC20^Y181G^ were incubated with 15 µM C18-Bz palmitate probe in DMEM (0.5% FBS) for 8 hours, after which the cells were lysed with the lysis buffer (HEPES (50 mM, pH 7.4), 0.5% NP-40, 0.25% SDS, 10 mM NaCl, 2 mM MgCl_2_, protease inhibitor cocktail (Roche), 0.20 µL/mL Benzonase). The click reaction was carried out using biotin-PEG-azide. The proteins were precipitated with chloroform/methanol, and pellets resuspended in 50 mM HEPES (pH 7.4 with 1% SDS, then diluted to 0.2%). Biotinylated proteins were enriched on Pierce™ NeutrAvidin™ Agarose beads for 2 hours. Following the washing steps, a solution containing triethanolamine (TEA, pH 7.5, 50 mM), 4 mM EDTA, and 0.5% ProteaseMAX was added. Then, 2.25 M neutralized hydroxylamine (HA) in 50 mM TEA (pH 7.5) with 4 mM EDTA was added and incubated for 2 hours. Chloroacetamide (CAA) was added to the supernatant at a final concentration of 15 mM, incubated for 15 minutes, and diluted with HEPES (50 mM, pH 8.0). The samples were then digested with 0.3-0.5 µg of trypsin at 37°C overnight, and prepared for MS analysis.

RAW files were uploaded into MaxQuant (version 1.6.11.0) and searched against Uniprot curated mouse proteome (as of 2019) using the built-in Andromeda search engine. Cysteine carbamidomethylation was selected as a fixed modification, and methionine oxidation and acetylation of protein N-terminus as variable modifications. Trypsin was set as the digestion enzyme, up to two missed cleavages were allowed and a false discovery rate of 0.01 was set for peptides, proteins and sites. Data was quantified using LFQ with a minimum ratio count = 2. Data analysis was performed using Perseus (version 1.6.2.1). MaxQuant proteingroups.txt output files were uploaded and filtered against contaminants, reverse and proteins identified by site and a base 2 logarithm was applied to all LFQ intensities. Missing values were imputed from a normal distribution (width = 0.3, downshift = 1.8) or by imputation of the lowest intensity for individual samples. A two-sample Student t-test was performed comparing WT ZDHHC20 with mutant Y181G ZDHHC20 (S0 = 0.1/0.5, FDR = 0.01/0.05) for all proteins remaining in the dataset and the results analysed according to their statistical significance.

### Cell surface protein biotinylation

Cell surface proteins were isolated by biotinylation with a cell-impermeable labelling reagent Sulfo-NHS-SS-biotin (Thermo), according to manufacturer’s instructions. The samples were then prepared for mass spectrometry on an Orbitrap Fusion Lumos Tribrid Mass Spectrometer (Thermo). All raw files were processed with MaxQuant v1.6.0.13 using standard settings and searched against the UniProt KB with the Andromeda search engine integrated into the MaxQuant software suite.

### Palmitoylation assay

HEK293T cells were transfected with a FLAG-tagged human DHHC20-expressing plasmid and a corresponding HA-tagged substrate-expressing plasmid using FuGENE transfection reagent (Promega) and incubated overnight. Alternatively, stable cell line expressing ZDHHC20^Y181G^ was used directly. Cells were treated with 20 µM alkyne-palmitate probe for 4 hours. Lysis was carried out using a lysis buffer (Tris 50 mM pH 7.4, 150 mM NaCl, 10% glycerol, 1% DDM, 0.5 mM TCEP, 10 µM palmostatin B, EDTA-free protease inhibitor cocktail (Roche)). Anti-HA beads were used for pull-down, followed by the TAMRA-azide click reaction (100 µM TAMRA-azide, 1 mM CuSO_4_, 1 mM TCEP, 100 µM TBTA, 1% NP-40 in PBS). The samples were then prepared for western blot and in-gel fluorescence. In-gel fluorescence was detected using Typhoon FLA 9500 scanner (GE Healthcare).

### Extracellular vesicle concentration and size analysis

Cells were plates at a density of 200,000 per well in a 6-well plate. After 24 h, the wells were washed three times with PBS, and exactly 2 mL of DMEM (0% FBS) added to the cells. After 24 h, the supernatant was collected, passed through a 0.45 µm filter, and analysed using Nanosight NS3000 instrument (Malvern Panalytical) according to the manufacturer’s instructions.

### Statistical analysis

Mead’s Resource Equation was used to predetermine sample size for intravenous injection mouse experiments: E = N – T – B, where E is the error degrees of freedom, N is the total degrees of freedom (i.e. the total number of experimental units (animals) minus one), T is the treatment degrees of freedom (i.e. the total number of experimental groups minus one), and B is the blocks degree of freedom (i.e. the total number of blocks (cages) minus one). The value of E was aimed to be between 10 and 20, as previously suggested, to balance the statistical power and resource utilisation (Festing M et al., 2002). The experimental unit was defined as an individual animal. Randomisation method used was block randomisation, where the blocking unit was a cage. The treatments were allocated randomly within each cage. Sample size was not predetermined for the KPCZ survival analysis, and the animals for this experiment were recruited on a rolling basis. Sample size for *in vitro* experiments was not predetermined, and these were carried out in biological triplicates. The exact sample size and statistical test for each dataset are reported for each figure. Blinding to the genotype was performed for any manual analysis of tissue histology. Data analysis was carried out using GraphPad Prism 9.4.0 software. Protein datasets were overlapped to create area-proportional Venn diagrams using BioVenn web application (Hulsen et al., 2008). The Kaplan-Meier survival curve was plotted using the RNA-seq data from The Cancer Genome Atlas (TCGA), extracted from the Human Protein Atlas survival analysis tool.

### Data and Software Availability

The accession numbers for the DNA/RNA sequencing data reported in this paper are GSE223623 and GSE220914. The mass spectrometry proteomics data have been deposited to the ProteomeXchange Consortium via the PRIDE (Perez-Riverol et al., 2022) partner repository with the dataset identifiers PXD036914, PXD038048, and PXD038384. Detailed protocols for selected experiments described in this paper are available to view and comment at protocols.io (https://www.protocols.io/workspaces/GT_protocols). A detailed list of reagents is provided in Supplementary Table 7. All source data and supplementary files are available on Figshare (https://figshare.com/s/204aeb6c53bbe81cacc9).

## Acknowledgments

The authors would like to thank the science technology platforms at the Francis Crick Institute: Biological Research Facility (Clare Watkins, Peter Owers), Experimental Histopathology (Emma Nye), Proteomics STP (Bram Snijders, Joanna Kirkpatrick, Mark Skehel, Helen Flynn), Bioinformatics and Biostatistics (Stuart Horswell, Stefan Boeing, Philip East), Advanced Sequencing Facility (Nik Matthews), and Cell Services. We thank David Tuveson for PDA530Met and FC1199 cell lines, and the members of the Tuveson group for their help (Lindsey Baker, Youngkyu Park, Ashley Adler, Erin Brosnan). We thank Miriam Molina Arcas and David Hancock for their help. This work was funded by CRUK/EPSRC Multidisciplinary Award to E.W.T. and J.D. (C29637/A27506 and NS/A000078/1), CRUK Programme Grant to E.W.T. (DRCN PG-Nov21\100001), Wellcome Trust Senior Investigator Award to J.D. (103799/Z/14/Z), and European Research Council Advanced Grant RASImmune to J.D. This work was supported by the Francis Crick Institute, which receives its core funding from Cancer Research UK (FC001070), the UK Medical Research Council (FC001070), and the Wellcome Trust (FC001070), awarded to J.D. G.T received Fulbright-CRUK Scholar award to conduct part of the research at Cold Spring Harbor Laboratory.

## Author Contributions

Conceptualisation, J.D., C.S., G.T., and E.W.T.; Methodology, G.T., C.S., and A.Y.R.; Validation, A.Y.R.; Formal Analysis, G.T. and C.S.; Investigation, G.T., C.S., A.Y.R., M.P.B., J.S., B.S., C.A.O., A.S-B., S.L.P., and E.H. Data Curation, G.T.; Writing – Original Draft, G.T. and C.S.; Writing – Review & Editing, all authors; Visualisation, G.T.; Supervision, J.D. and E.W.T.; Project Administration, J.D. and E.W.T.; Funding Acquisition, J.D. and E.W.T.

## Rights and Permissions

For the purpose of Open Access, the author has applied a CC BY public copyright licence to any Author Accepted Manuscript version arising from this submission.

## Supplementary figures

**Figure S1.**
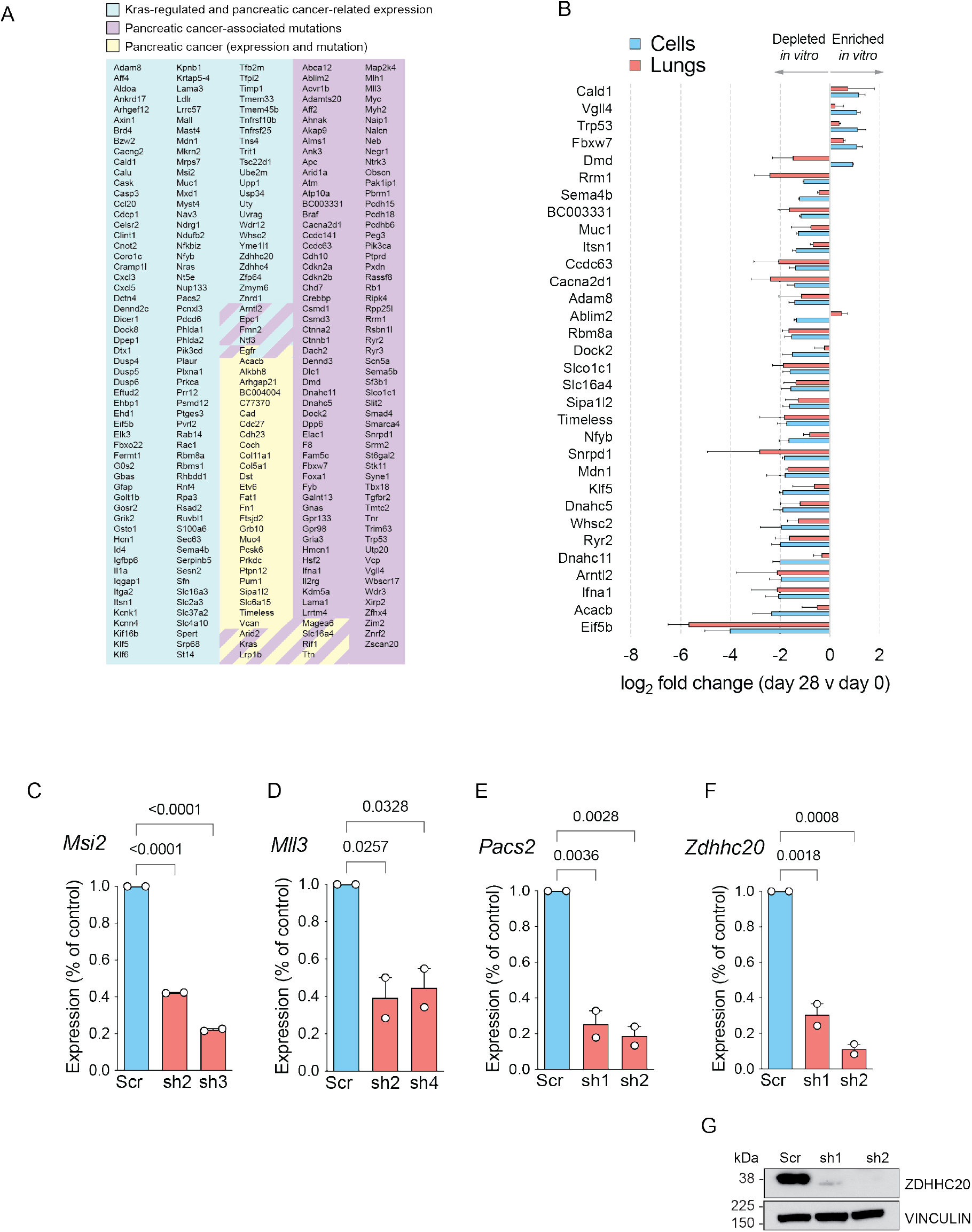
ShRNA screen identifies different dependencies of pancreatic cancer-relevant genes *in vitro* versus *in vivo*. Related to Figure 1. **(A)** Categories of genes targeted by shRNAs in the screen. **(B)** shRNAs hits *in vitro* and any associated enrichment or depletion *in vivo* (average log_2_ fold change of multiple shRNAs per gene). **(C-F)** Validation of gene expression knock-down following transduction with shRNAs targeting *Msi2* **(C)**, *Mll3* **(D)**, *Pacs2* **(E)**, and *Zdhhc20* **(F). (G)** Western blot image of ZDHHC20 protein level in shRNA-treated cells.

**Figure S2.**
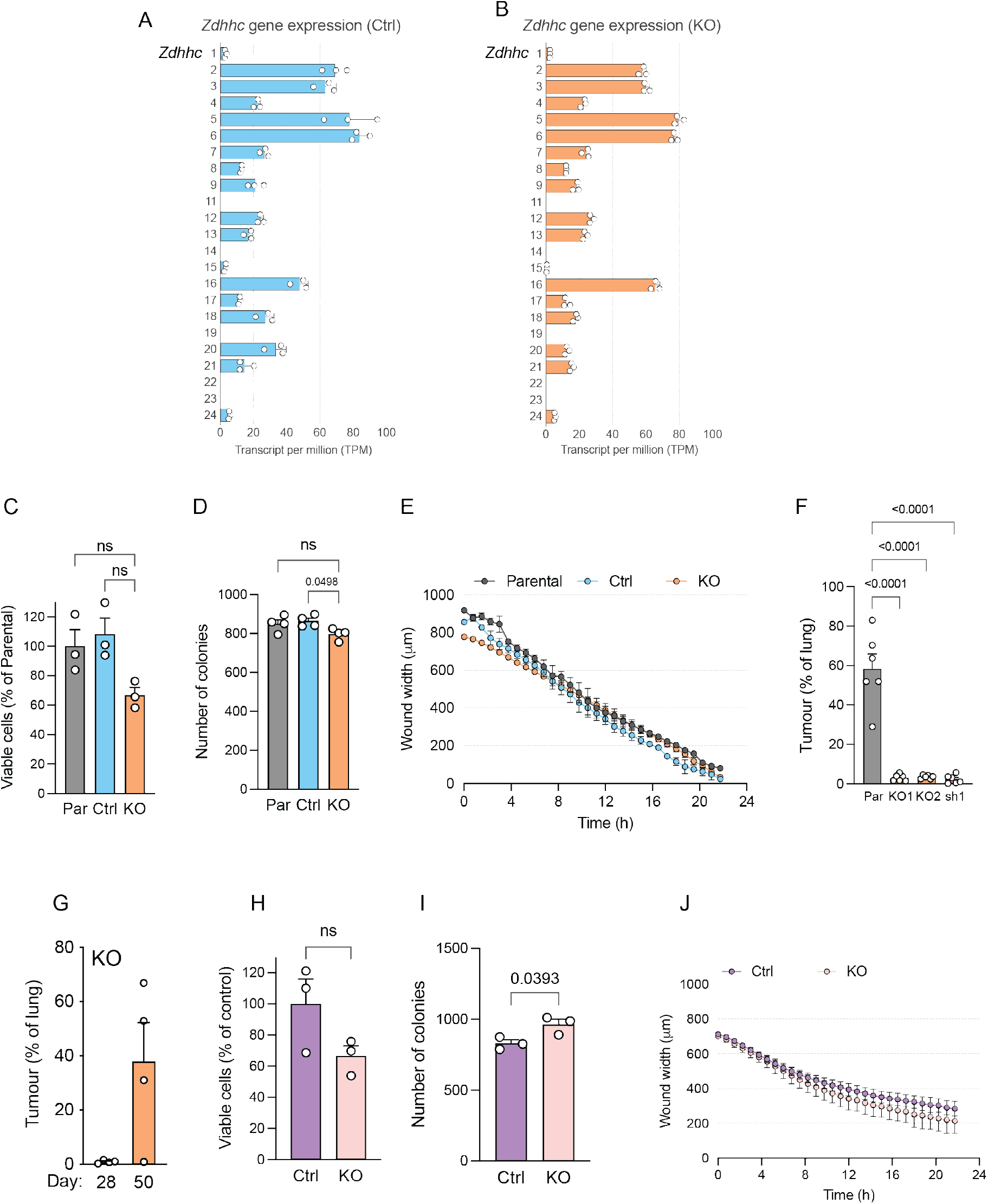
Characterisation of Zdhhc20 KO cells. Related to Figure 2. **(A-C)** Gene expression of the *Zdhhc* family members in PDA530Met control cells **(A)**, *Zdhhc20* KO **(B)**, and their log_2_ fold change in KO versus Ctrl cells **(C). (D)** Anchorage-independent growth of Zdhhc20 KO clones of PDA530Met cells in 3D culture. **(E)** Quantification of colonies of *Zdhhc20* KO clones of PDA530Met cells in 3D Matrigel culture. **(F)** Migratory capacity of *Zdhhc20* KO cells in the scratch-wound assay (n=2). **(G)** Quantification of tumour area in the lungs of mice injected with parental cells, *Zdhhc20* shRNA KD cells, and two independent *Zdhhc20* KO clones. **(I)** Quantification of tumour area in the lungs at day 28 and 50 after i.v. injection of *Zdhhc20* KO PDA530Met cells. **(J)** Anchorage-independent growth of FC1199 control and *Zdhhc20* KO cells in 3D culture. **(K)** Analysis of colony frequency of FC1199 control and *Zdhhc20* KO clones in 3D Matrigel culture. **(L)** Migration of FC1199 control and *Zdhhc20* KO cells in the scratch-wound assay (n=2). Data are shown as mean + S.E.M. Ordinary one-way ANOVA with Tukey’s multiple comparisons test was used.

**Figure S3.**
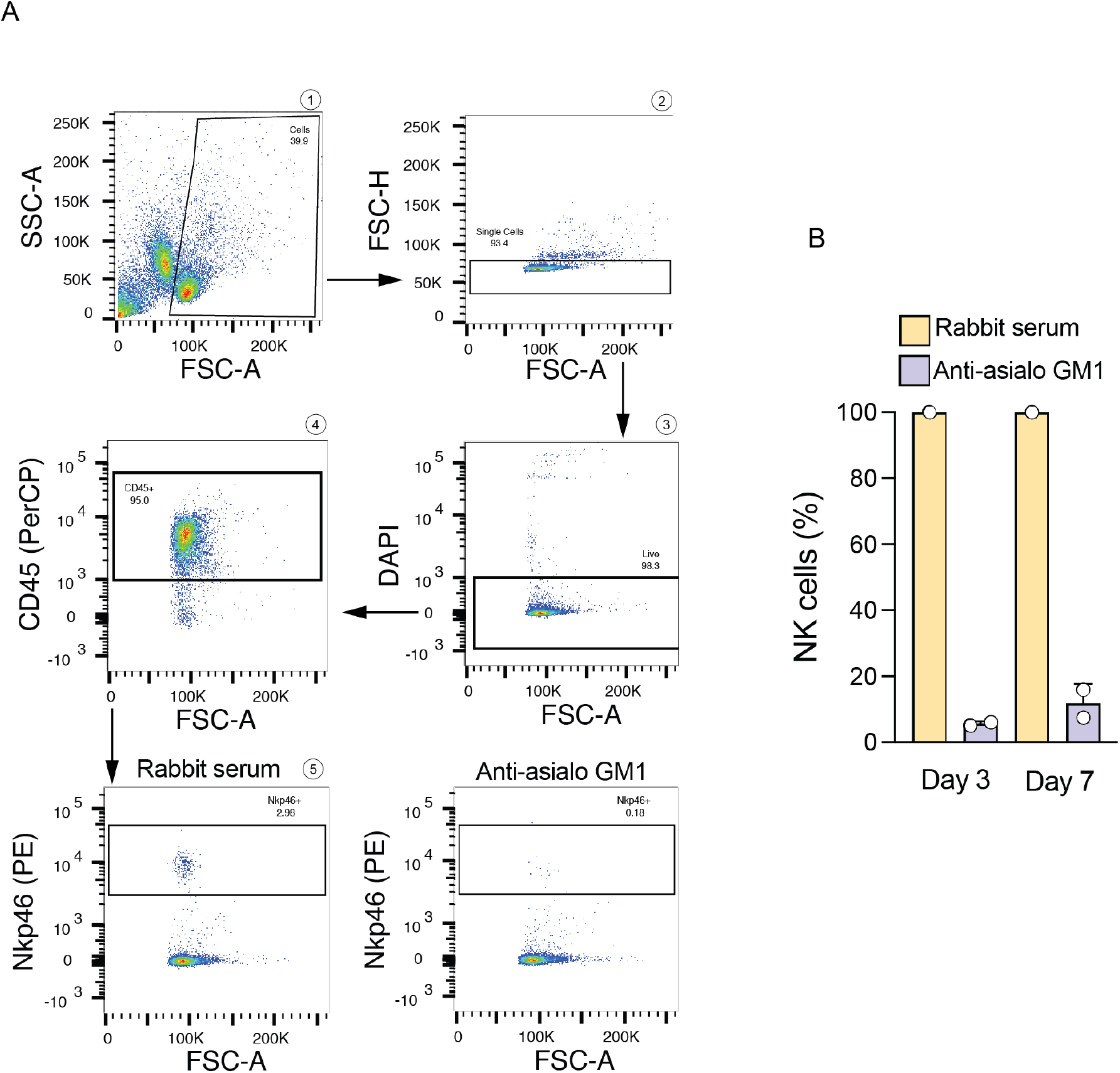
Characterisation of NK cell depletion. Related to Figure 3. **(A)** FACS diagrams showing the gating strategy for quantifying the number of NK cells in mouse spleen. **(B)** Depletion of NK cells in wild type 129S mice following administration of the anti-asialo GM1 serum.

**Figure S4.**
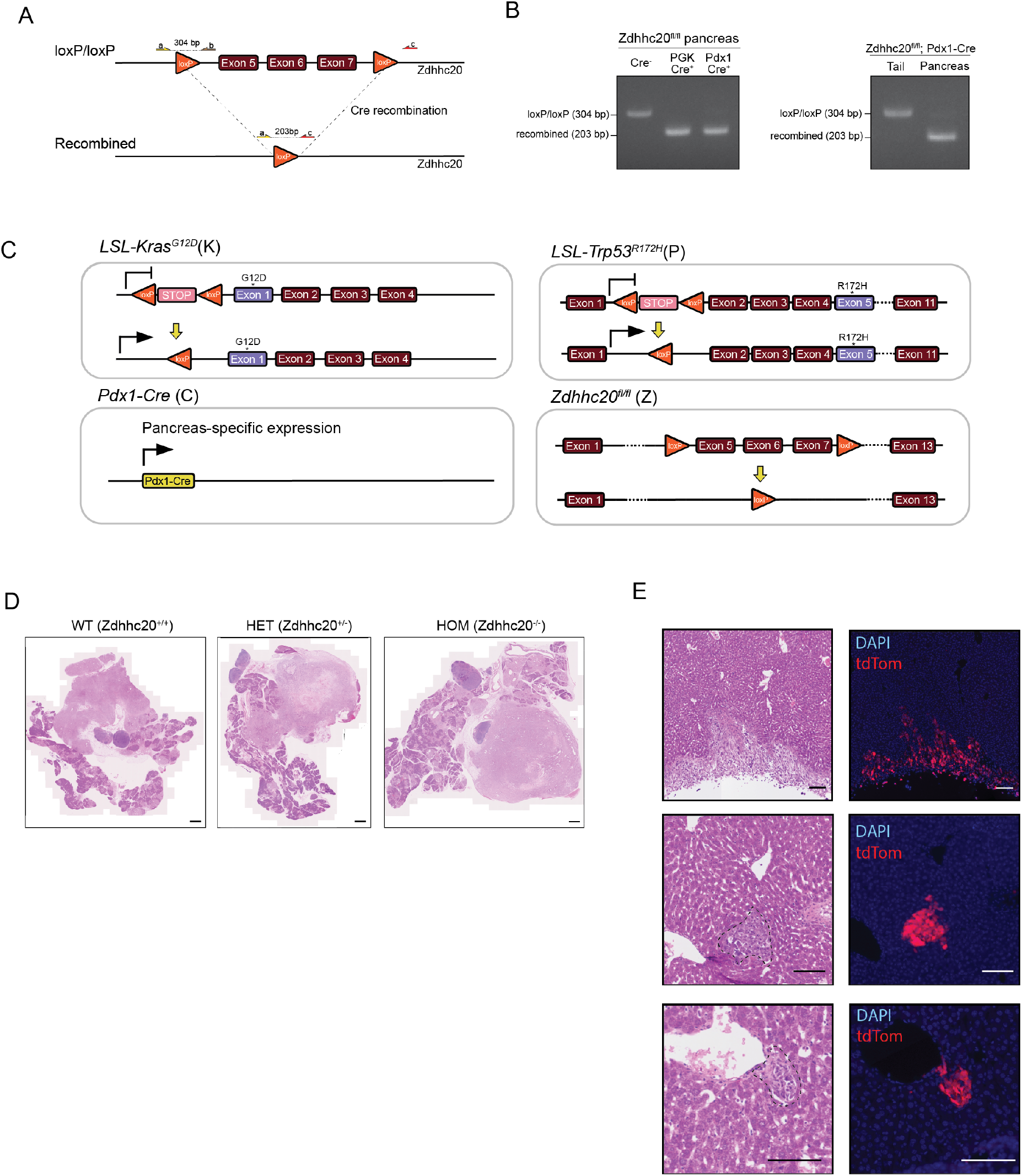
Characterisation of *Zdhhc20* KO mouse model. Related to Figure 4. **(A)** Schematic of genotyping strategy to confirm successful Cre recombination. **(B)** Agarose gel electrophoresis images showing complete and pancreas-specific recombination of floxed *Zdhhc20* allele. **(C)** Schematic of the KPCZ mouse model. **(D)** Representative images of H&E sections of primary tumours in KPCZ mice, Scale bar, 1000 µm. **(E)** Representative images of H&E sections and corresponding fluorescence images of selected microscopic lesions in the liver to validate the scored lesions are of pancreatic origin. Scale bar, 100 µm.

**Figure S5.**
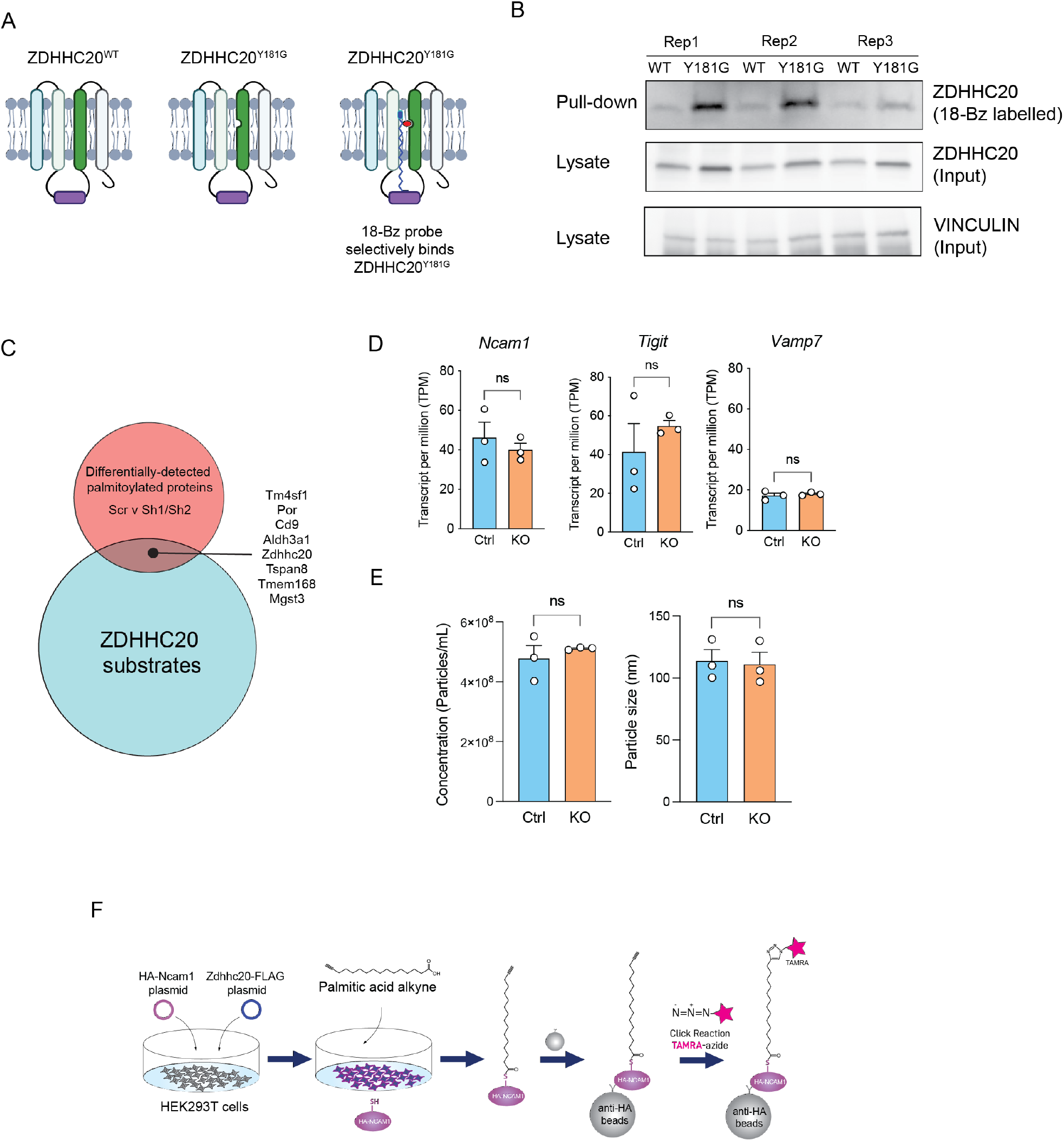
Validation of ZDHHC20 substrate profiling method. Related to Figure 4. **(A)** Schematic of Y181G ZDHHC20 mutant and chemical probe binding. **(B)** Loading assay of WT and Y181G ZDHHC20. **(C)** Overlap between ZDHHC20 substrates and differentially-palmitoylated proteins in shRNA cells. **(D)** Gene expression of *Ncam1, Tigit*, and *Vamp7* in Ctrl and KO cells as determined by RNAseq. **(E)** Comparison of size and number of extracellular vesicles in Ctrl and KO cells. **(F)** Schematic of the palmitoylation assay used to generate the data in Fig5H.

